# The structure of an Amyloid Precursor Protein/talin complex indicates a mechanical basis of Alzheimer’s Disease

**DOI:** 10.1101/2024.03.04.583202

**Authors:** Charles Ellis, Natasha L Ward, Matthew Rice, Neil J Ball, Pauline Walle, Chloé Najdek, Devrim Kilinc, Jean-Charles Lambert, Julien Chapuis, Benjamin T Goult

## Abstract

Misprocessing of Amyloid Precursor Protein (APP) is one of the major causes of Alzheimer’s disease. APP is a transmembrane protein comprising a large extracellular region, a single transmembrane helix and a short cytoplasmic tail containing an NPxY motif (normally referred to as the YENPTY motif). Talins are synaptic scaffold proteins that connect the cytoskeletal machinery to the plasma membrane via binding to one of two highly conserved NPxY motifs in the cytoplasmic tail of integrin transmembrane receptors. Here we report the crystal structure of an APP/talin1 complex identifying a new way to couple the cytoskeletal machinery to synaptic sites via APP. Proximity Ligation Assay (PLA) confirmed the close proximity of talin1 and APP in primary neurons, and we show that talin1 depletion has a dramatic effect on APP processing in cells. Structural modelling indicates that APP has the capacity to form an extracellular meshwork that mechanically couples the cytoskeletal meshworks of both the pre-, and post-synaptic compartments. In this context, we propose APP processing as a mechanical signalling pathway with similarities to Notch signalling, whereby the cleavage sites in APP represent mechanical sensors, with varying accessibility to cleavage by secretases. During synaptogenesis in healthy neurons, the APP/talin linkage would provide an exquisite mechanical coupling between synapses, with tightly controlled APP processing providing instructions to maintain this synchrony. Furthermore, APP directly coupling to the binary switches in talin indicates a role for APP in mechanical memory storage as postulated by the MeshCODE theory. The implication that APP is a regulator of mechanical signalling leads to a new hypothesis for Alzheimer’s disease, where mis-regulation of APP dynamics results in loss of mechanical integrity of the synapse, corruption and loss of mechanical binary data, and excessive generation of the toxic plaque-forming Aβ42 peptide. Much needs to be done to experimentally validate this idea, but we present here a novel theory of Alzheimer’s Disease with a role for APP in the mechanically coded binary information storage in the synapse, which identifies a potential novel therapeutic strategy for treating Alzheimer’s Disease.

**Graphical Abstract.**
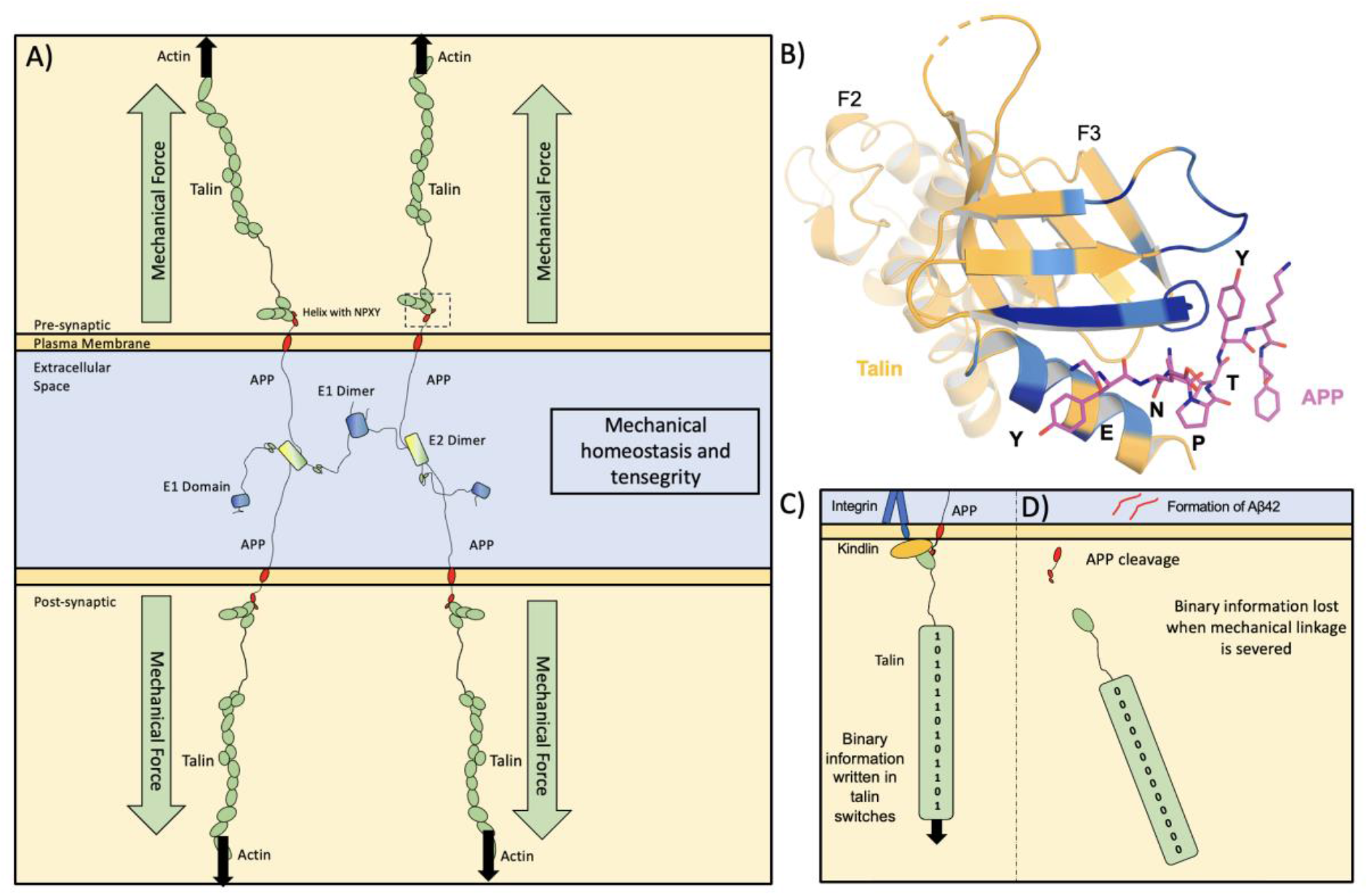
Graphical Abstract Legend: Amyloid Precursor Protein (APP) forms an extracellular meshwork that provides a coupling between the mechanical signalling machinery of the pre– and post-synaptic neurons. **A**) APP binds to talin indicating that APP will experience mechanical forces which leads us to propose that APP processing is a Notch-like signalling pathway that maintains mechanical homeostasis at the synapse. **B)** The crystal structure of APP bound to talin F2F3. **C)** APP is part of the adhesion complex that attaches talin to the membrane. **D)** The loss of adhesion integrity in the synapse, or other factors that lead to misprocessing of APP will perturb this coupling and contribute to Alzheimer’s Disease. See also <u>Movie 1</u>.

## Introduction

Alzheimer’s disease (AD) is a leading cause of dementia, accounting for 60-80% of total dementia cases (DeTure and Dickson, 2019). The worldwide cost of AD has been estimated to be ∼$604 billion per year (Langa, 2015). AD is characterised by the presence of amyloid plaques and tau tangles (DeTure and Dickson, 2019), which are taken into account in the National Institute of Aging and Alzheimer’s guidelines for AD patient pathophysiology (McKhann et al., 2011; Montine et al., 2016), including measures such as Braak neurofibrillary tangle stage (Braak et al., 2006; Braak and Braak, 1991; Braak and Braak, 1997), Thal phases of amyloid deposition (Thal et al., 2002) and the Consortium to Establish a Registry for Alzheimer Disease (CERAD) score of neuritic amyloid plaques (Mirra et al., 1991). The development of amyloid plaques occurs as a result of defects in the processing of Amyloid Precursor Protein (APP) (Selkoe, 1998). APP is expressed in most tissues but is most highly expressed in the brain (Karlsson et al., 2021). Surprisingly, despite the extensive research effort to understand APP biology, the cellular roles of APP remain unclear, in part due to the shift in focus to the toxic species that forms the amyloid plaques in AD, amyloid-β (Aβ42), a 42-residue cleavage product of APP (Findeis, 2007; Sengupta et al., 2016). While APP’s roles are not yet fully understood, several functions have been identified including cell-cell adhesion, neurite outgrowth and synaptogenesis, cell migration, cell signalling and apoptosis (Zheng and Koo, 2006).

## Synaptic adhesion and mechanical signalling

Synapses represent the perfect system for the discrete transfer of information between two cells. To enable precision in this information transfer, the synaptic boutons are connected by a complex network of scaffolding molecules. This scaffolding is anchored in two ways, by cell-cell adhesions, and via connections between the cell and the surrounding meshwork of proteins called the Extracellular Matrix (ECM) (Levy et al., 2014; Wlodarczyk et al., 2011). Many synaptic cell adhesion molecules exist that connect the two synaptic boutons to the ECM. These adhesion complexes are coupled to the intracellular cytoskeletal machinery via adapter proteins and large signalling complexes assemble on these linkages. In synapses these cell-cell and cell-matrix couplings are incredibly complex, involving many proteins that ensure a perfectly tuned connection for efficient neuronal signalling (Dalva et al., 2007; Missler et al., 2012).

## The MeshCODE theory and the role of talin in mechanical signalling

One major family of ECM receptors are the integrins (Hynes, 1992), which are essential for the formation of synapses (Huang et al., 2006; Kerrisk et al., 2013; Ning et al., 2013; Webb et al., 2007). Large mechanical signalling machinery, known as integrin adhesion complexes, assemble on the short cytoplasmic tail of integrins. Stable adhesions require cytoskeletal connectivity and provide a mechanosensitive coupling that instructs the cell how to behave in the context of the local environment. The central adapter that connects integrins to the actin cytoskeleton is the protein talin, which contains 13 helical bundles that behave as force-dependent binary switch domains (Goult et al., 2013; Yao et al., 2016). Each switch can be converted between folded “0” and unfolded “1” states by small changes in contractility that can be generated by the cell’s motor proteins. The presence of binary switches in the synaptic scaffolds led to the idea that information can be written into the switch patterns of the scaffolds, representing a meshwork code, described in the MeshCODE theory (Goult, 2021; Goult et al., 2021). In this theory, these switch patterns represent a binary code the cell uses to spatially organise the enzymatic processes in the synapse to coordinate synaptic activity (Ball et al., 2024; Barnett and Goult, 2022). Currently, the role of talin in synaptic regulation and memory is not well studied, however, the role of talin in mechanical memory in other systems has been demonstrated experimentally (Dahal et al., 2022; Marhuenda et al., 2023).

## APP and integrin adhesion complexes

APP co-localises with integrins and loss of APP has been shown to result in decreased β1 and β3-integrin expression (Pfundstein et al., 2022; Ristori et al., 2020; Yamazaki et al., 1997). APP has also been shown to colocalise with talin (Storey et al., 1996) and talin2 has been implicated in AD previously (Gusareva et al., 2018). Furthermore, mechanotransduction downstream of integrins has also been linked to AD (Donnaloja et al., 2023). More recently, genome-wide association studies (GWAS) cast focal adhesion (FA) proteins in a central role in AD pathology (Dourlen et al., 2019; Eysert et al., 2021), and knock down of FA proteins significantly perturbs APP processing (Chapuis et al., 2017). Kindlin2 (FERMT2), a key co-regulator of integrin activity with talin, was shown to be a major modulator of APP metabolism (Chapuis et al., 2017).

APP is an integral membrane protein that contains a large extracellular region comprised of an N-terminal E1 domain, a Kunitz-type protease inhibitor (KPI) domain, an E2 domain, connected to a single transmembrane helix (TMH) (Fig.1). The linker length between the E2 and TMH is invariant in length in all isoforms, ∼34 nm (Fig.1B). Intracellularly, APP is comprised of a short cytoplasmic tail, sometimes referred to as the APP intracellular domain (AICD), which contains a completely conserved NPxY motif (Chen et al., 1990), more commonly referred to as the YENPTY motif. APP is shown drawn to scale in Fig.1D,E, we show isoform APP770 as it contains all the APP domains, however APP695, the alternatively spliced, neuronal specific isoform, lacks the KPI domain (Kang and Müller-Hill, 1990; Rohan de Silva et al., 1997).

**Figure 1.**
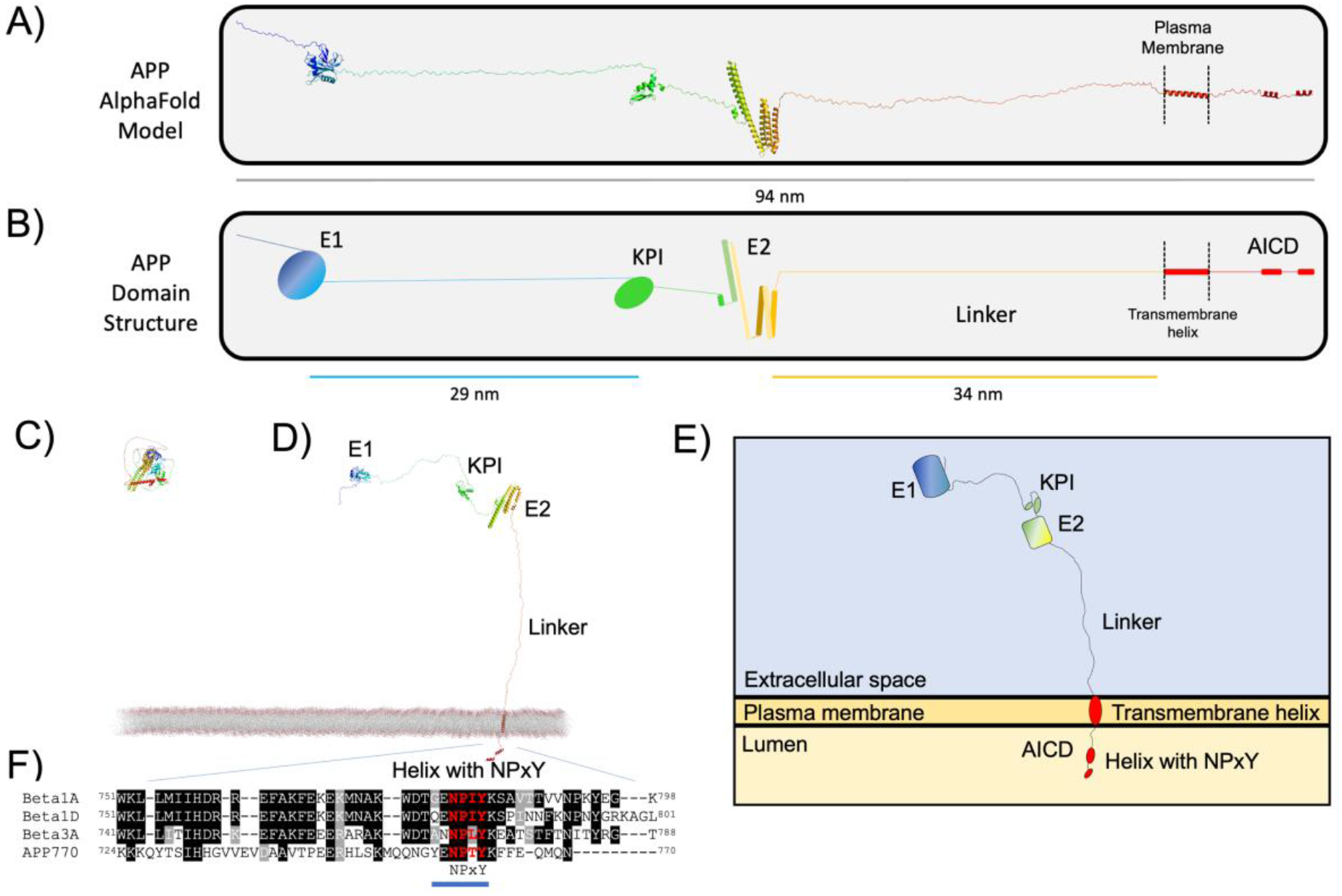
Amyloid Precursor Protein (APP) to scale. **A-B**) APP is an integral membrane bound protein, comprising a large extracellular region, a single transmembrane helix and a short cytoplasmic tail referred to as the AICD, that contains a highly conserved NPxY motif. The largest isoform, APP770 is shown. **A)** The unfurled AlphaFold structural model showing the structured domains in the context of the entire molecule. **B)** Cartoon representation of A), the colour coding for the domains is used in all the schematics. The scale bars show the lengths of the linker regions. **C-E)** APP in the context of the membrane. **C)** The AlphaFold structural model of APP. **D)** The unfurled AlphaFold model from C) shown embedded in the plasma membrane, showing the dimensions of the protein and arrangement of the domains relative to each other and the membrane. **E)** Cartoon representation of D). **F)** Sequence alignment of the human cytoplasmic tails of APP and integrins β1A, β1D and β3. The NPxY-containing YENPTY motif is shown by a blue line.

The presence of an NPxY motif in APP, coupled with the published data on the role of integrin adhesion complexes in APP processing (Dourlen et al., 2019; Eysert et al., 2021) prompted us to analyse whether talin and APP might interact. In our previous work on the role of talin in the synapse we undertook an in-depth structural analysis of the talin protein and its interactions, drawing these molecules “to scale” (Barnett and Goult, 2022). Here, we carried out a similar structural analysis of APP and its interactions. We harnessed the published structural data on APP in the Protein Database and coupled it to the structural models in the AlphaFold database (Jumper et al., 2021; Varadi et al., 2021). This analysis, coupled with the crystal structure of the APP NPxY motif bound to talin, leads us to propose a new role for APP as a mechanocoupler, connecting the cytoskeletal force generation machinery of the two sides of the synapse. This leads to a novel hypothesis for a mechanical basis of APP processing, and a new theory for memory loss in AD, caused by the loss of binary information written into the shapes of the talin molecules scaffolding each synapse as the mechanical homeostasis of the synapse is lost. As such, the scientific framework outlined, provides a new direction for AD research and treatment.

## Results

### The NPxY motif in the cytoplasmic tail of APP binds to talin

The presence of an NPxY motif in the cytoplasmic tail of APP indicated that it would bind to FERM domain containing proteins. Sequence alignment of the NPxY motif of APP with those from β1A, β1D and β3 integrin confirmed the sequence homology that also extends to the membrane proximal part of the cytoplasmic tail (Fig.1F).

Talin is a large and complex protein comprised of 18 domains (Fig.2A,B) with the F3 subdomain of the talin head binding to the NPxY motif of the β-integrin subunit (Anthis et al., 2009; Calderwood et al., 2002). To test whether the APP NPxY motif might also interact with talin F3 we used Nuclear Magnetic Resonance (NMR) where ^15^N-labelled talin F3 was titrated with synthetic peptides of the APP cytoplasmic tail (residues 732-770), either wildtype or a mutated version where the NPTY residues were replaced with four alanine residues, hereafter called APP(4A) peptide. Addition of wildtype APP peptide resulted in chemical shift changes in the spectra of F3 of both talin1 (Fig.2C and Supp. Fig.1) and talin2 (Fig.2D and Supp. Fig.2), which mapped onto the β-integrin binding site (Fig.2I) confirming that the APP peptide binds in the NPxY-binding pocket on talin F3, as expected. Minimal chemical shift changes were observed on addition of the APP(4A) peptide confirming that the interaction is NPxY-dependent (Fig.2E,F).

**Figure 2.**
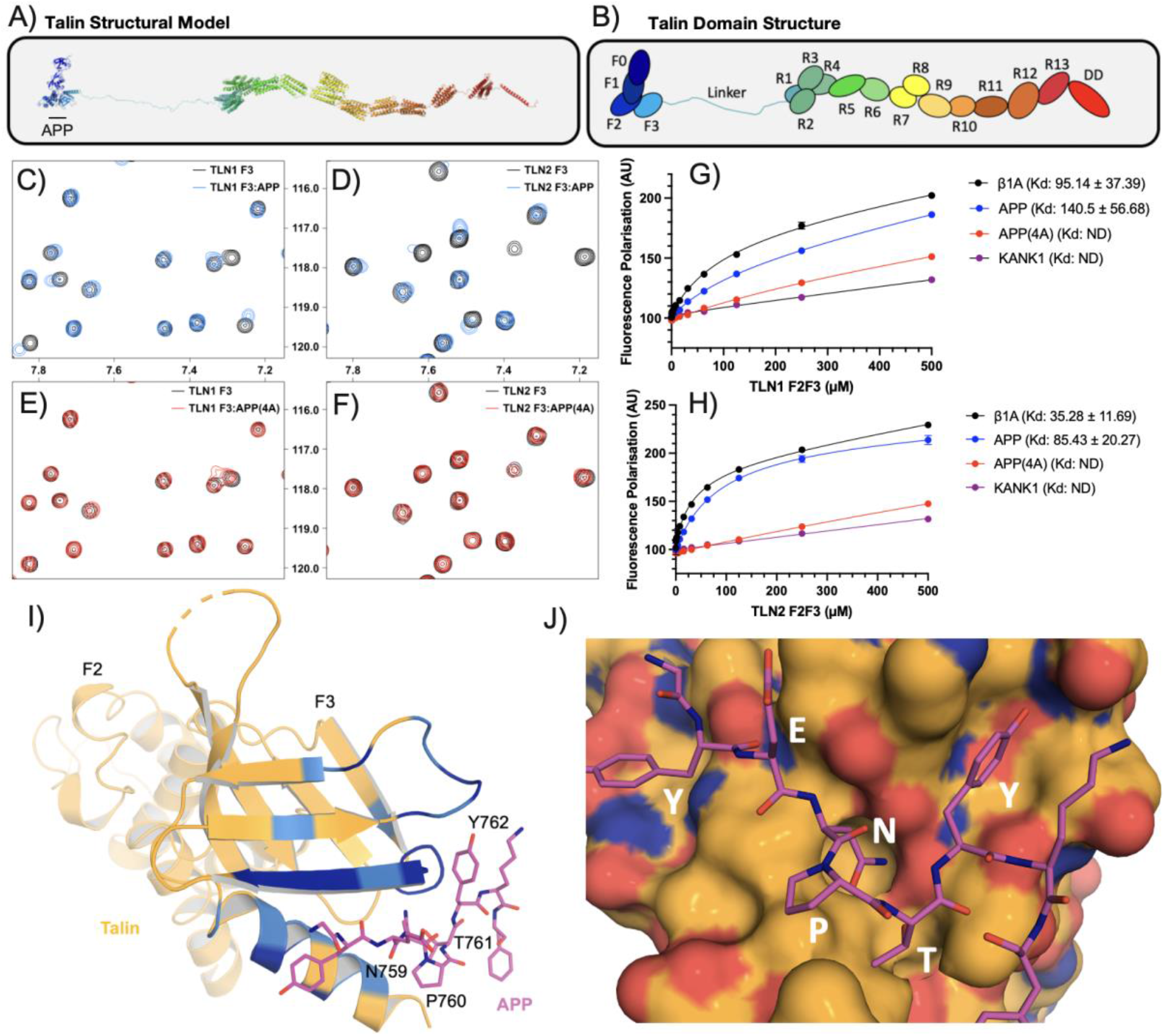
APP binds to talin directly. **A-B**) Talin is a large, synaptic scaffold molecule comprised of an N-terminal FERM domain and a large rod domain comprised of 13 helical bundles, R1-R13, that act as force-dependent binary switches. The F3 domain is the NPxY motif binding domain. **C-D)** ^1^H-^15^N HSQC spectra of ^15^N-labelled **C)** Talin1 F3 domain and **D)** Talin2 F3 domain in the absence (*black*) and presence of APP peptide (*blue*) at a ratio of 1:5. **E-F)** ^1^H-^15^N HSQC spectra of ^15^N-labelled **E)** Talin1 F3 domain and **F)** Talin2 F3 domain in the absence (*black*) and presence or APP(4A) peptide (*red*) at a ratio of 1:5. **G-H)** A fluorescence polarization assay was used to determine the K_d_ of the interaction between integrin β1A, APP, APP(4A) and **G)** talin1 and **H)** talin2. Dissociation constants ± SE (µM) for the interactions are indicated in the legend. All measurements were performed in triplicate. ND, not determined. **I)** The talin1/APP crystal structure. One heterodimer from the crystal structure of talin1 F2F3 bound to the APP cytoplasmic tail. The NPxY residues are labelled, and the colour coding is the chemical shift mapping from the NMR data in B). **J)** Surface representation of F3 coloured by electrostatics, showing the ^757^YENPTY^762^ motif bound. APP residues N759, T761 and Y762 fit into pockets on F3.

We next measured the binding affinity of APP to talin using a fluorescence polarisation assay. In this assay the APP and APP(4A) peptides are fluorescently tagged by covalently coupling a fluorescein-maleimide to a non-native terminal cysteine and titrated with both talin1 and talin2 F2F3 domains. This confirmed that i) both talin isoforms bind to APP and ii) the interaction is specific but weak (Fig.2G,H) as expected since talin also binds weakly to the NPxY motif of the β-integrin cytoplasmic tail in solution (with K_d_ in the range of 100-1000 µM) (Anthis et al., 2009; Calderwood et al., 2002; Garcıá –Alvarez et al., 2003). However, the interaction between talin and integrin is enhanced more than 1000-fold by the presence of negatively charged phospholipids (Moore et al., 2012) due to the tripartite interaction between talin, integrin and negatively charged phospholipid headgroups. Similarly, it is likely that the interactions between the positively charged residues on the talin head domain and negatively charged phospholipids also enhance the interaction of APP and talin.

### The crystal structure of talin1 F2F3 bound to the APP cytoplasmic domain

Early structural insights into the talin-integrin interaction came from crystal structures of a chimeric fusion of the talin F2F3 domains and the β3-integrin tail (Garcıá –Alvarez et al., 2003). Using the design strategy of the β3(739-749)-talin(209-400) chimera (Garcıá –Alvarez et al., 2003) we generated an APP(754-764)-talin(209-400) chimera, hereafter called the APP-talin chimera (Supp. Fig.3A). The APP-talin chimera crystals were birefringent flat plates (Supp. Fig.3B) belonging to space group P2_1_2_1_2_1_ which diffracted to 2.7 Å. The structure of the APP-talin chimera was solved by molecular replacement using the F2F3 domains (1MIX; (Garcıá – Alvarez et al., 2003)) as a search model with a single copy in the asymmetric unit. In contrast to the β3-integrin-talin chimera, which adopted a homodimeric arrangement in the asymmetric unit mediated by reciprocal F3-NPxY interactions of the two fusion proteins, the APP-talin chimera showed a “daisy chain” style arrangement where each APP tail interacted with the F3 domain of a symmetry related molecule (Supp. Fig.3C). The NPTY was well resolved in the density (Supp. Fig.3D) and closer inspection confirmed that the NPxY motif of the APP was binding canonically in the NPxY motif binding site on talin F3 (Fig.2I,J). The chemical shift changes upon addition of APP peptide in the NMR data precisely map onto the APP-interacting surface on F3 (Fig.2I).

### Structural modelling of APP as a synaptic cell adhesion molecule

The identification of a direct coupling between APP and the cytoskeleton, mediated via interaction with talin, supported an adhesion role for APP. Therefore, we next sought to explore the role of APP as a synaptic cell adhesion molecule. To do this we combined the known atomic structures of APP domains (Baumkötter et al., 2014; Dahms et al., 2010; Wang and Ha, 2004) with the AlphaFold model of the full-length APP to visualise the unstructured elements. As is visible in Fig.1 much of APP is unstructured, with intrinsically disordered regions connecting the folded globular domains. Both extracellular domains 1 and 2 (E1 and E2) have been shown to homodimerize, and crystal structures of the E1 homodimer (Fig.3B, PDB ID: 4JFN (Baumkötter et al., 2014) and PDB ID: 3KTM (Dahms et al., 2010)), and E2 homodimer (Fig.3C, PDB ID: 3NYL (Wang and Ha, 2004)) have been solved. We used these known dimerisation structures to superimpose four full-length APP structural models so as to orientate them based on the structural constraints of dimerisation. This combination of structural data with structural modelling immediately indicated a structural basis for how cis-(Supp Fig.4A,B) and trans-APP-mediated (Supp Fig.4C,D) couplings would work together to enable APP to serve as a synaptic adhesion molecule connecting the pre– and post-synaptic membranes (Fig.3).

**Figure 3.**
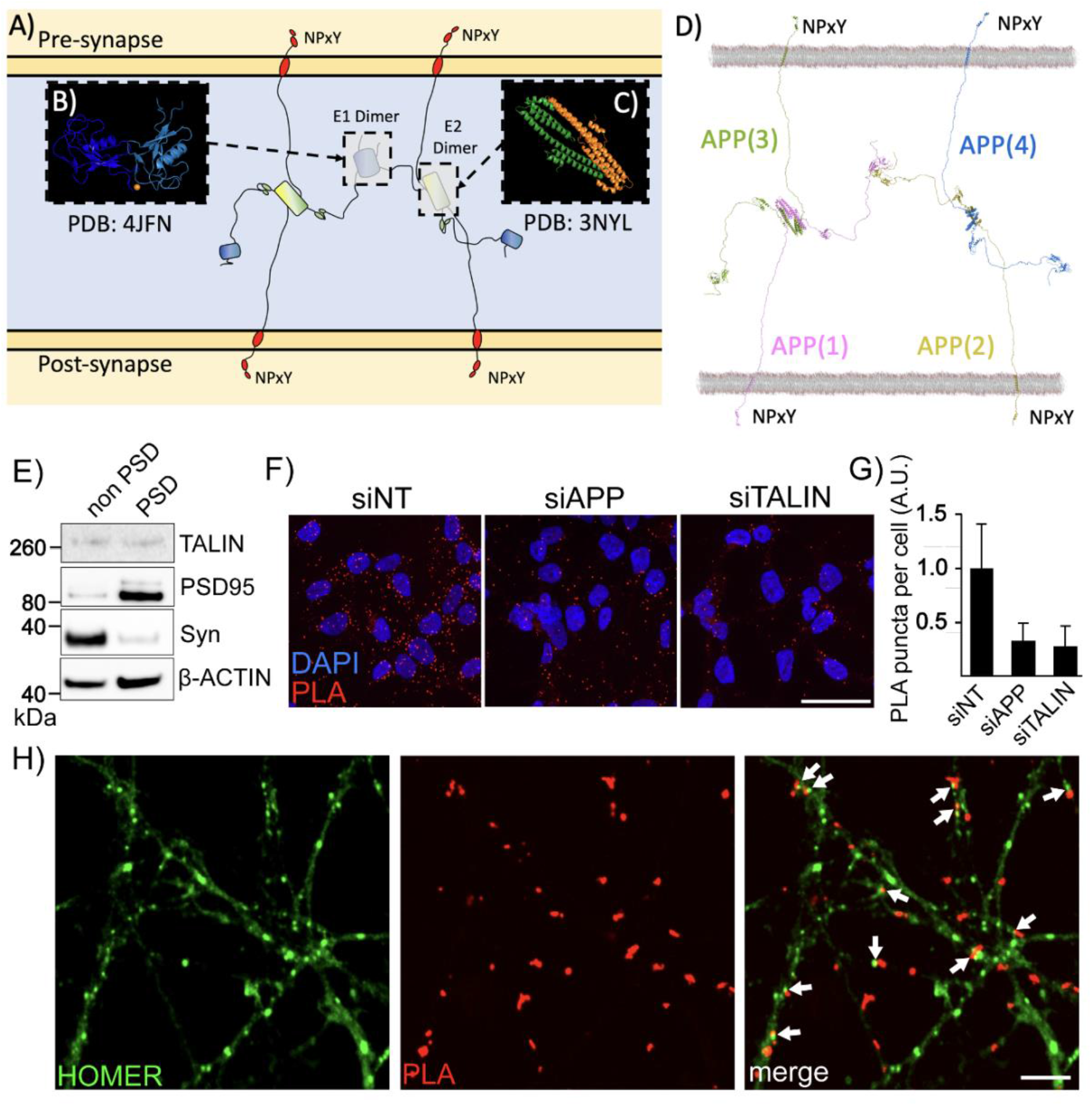
APP dimerisation leads to the formation of an extracellular synaptic meshwork. **A**) Four APP molecules are shown forming a dimer of dimers. **B-C)** The crystal structures of **B)** the E1 dimer (Baumkötter et al., 2014) and **C)** the E2 dimer (Wang and Ha, 2004). These dimer interfaces were used to overlay four full-length APP structural models. **D)** Structural model of the APP synaptic meshwork. The four APP molecules are numbered, 1-4, with APP molecules 1 (pink) and 2 (yellow) embedded in the post-synaptic membrane, and 3 (green) and 4 (blue) embedded in the pre-synaptic membrane. On the cytoplasmic face of both synaptic compartments are positioned NPxY motifs that are spatially organised by the APP oligomerisation. See also <u>Movie 2</u>. **E)** Synaptic fractionation experiment revealed the presence of talin1 in both pre– and post-synaptic compartments. Post Synaptic Density (PSD). **F-H) Proximity Ligation Assay (PLA) of APP and talin1 in cells and neurons. F)** PLA of APP/talin (red) signal in HEK293-APP^695WT^ transfected with siRNAs targetting APP or talin. A non-targetting (NT) siRNA was used as a control. The nucleus is visualised using Hoechst (blue). Scale bar = 40 µm. **G)** Quantification of PLA as performed in F). **H)** APP/talin PLA puncta (red) were also observed at synapses from primary neuronal cultures. Homer staining (green) was used as a synaptic marker. Scale bar = 4 µm.

**Figure 4.**
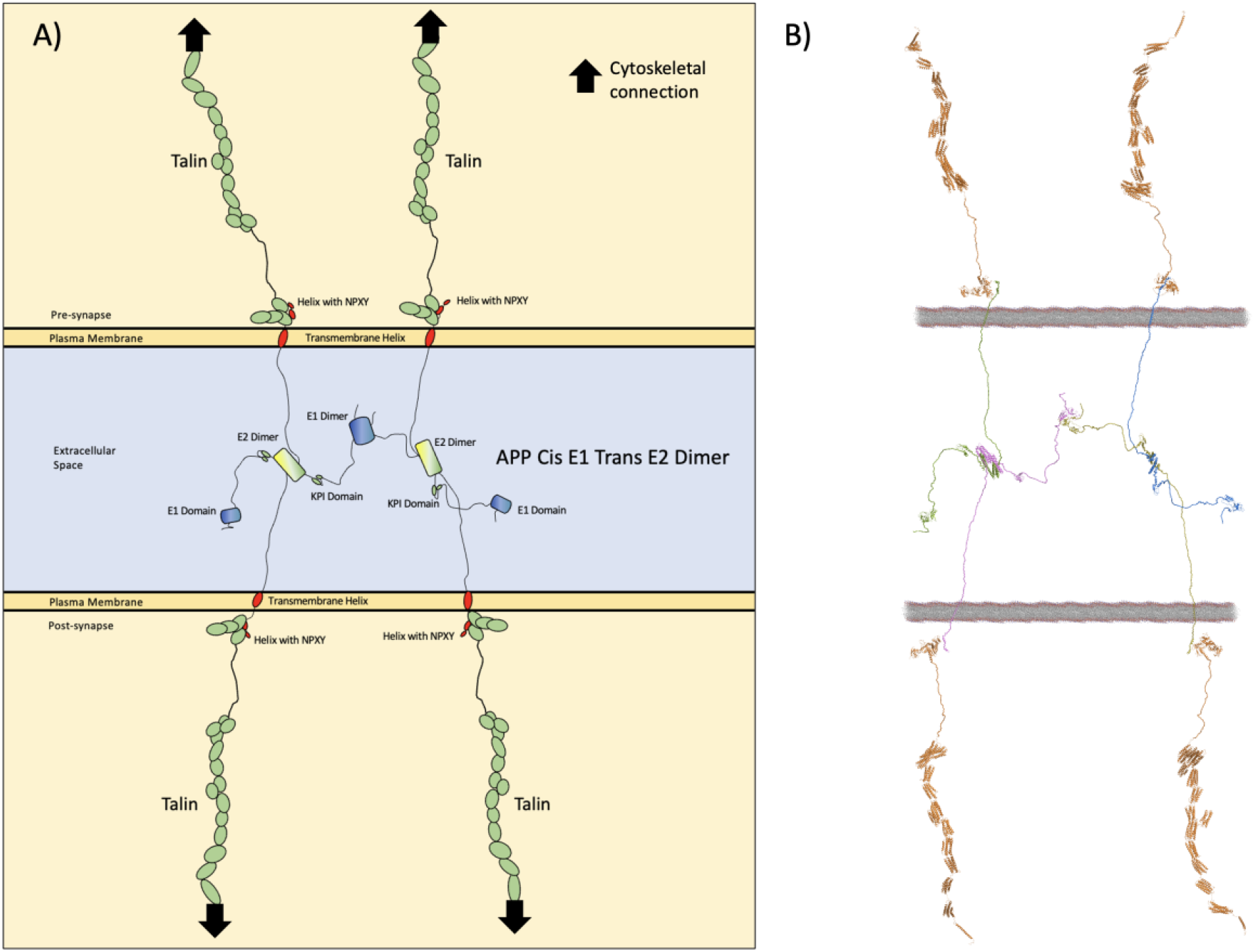
APP as a mechanocoupler coupling the two cells force-generation machineries via the talin mechanically operated signalling scaffolds. **A**) Cartoon representation and **B)** Structural model of four APP molecules, two from each side of the synapse forming a meshwork that spatially positions four talin binding sites, that can then recruit four talin molecules to connect the synaptic junction to the actin and microtubule cytoskeletons. The coupling to the actin cytoskeleton via talin is shown by black arrows.

What is striking about this arrangement is that it represents an extracellular protein meshwork, composed of APP, connecting the adjoining cells across the synaptic cleft. This arrangement resembles a hybrid of an ECM meshwork and a cell-cell junction, but one that is built by the two cells’ surface receptors. In essence, APP forms a hybrid cell-cell/cell-ECM coupling between cells. We note that the tetrameric APP arrangement shown in Fig.3 also indicates a mechanism for the formation of higher order species via the flailing, undimerised E1 domains so further oligomerisation of APP will likely occur forming a substantial meshwork between cells.

### Talin is in close proximity to APP in synapses

Next, we sought to investigate the APP/talin interaction in cellular models. First, by performing synaptosomal fractionations from primary neuronal cultures, we confirmed the presence of talin in both the pre– and post-synaptic compartments (**Fig. 3E).** Next, we developed a proximity ligation assay (PLA) able to detect potential APP/talin complexes using antibodies that recognise APP and talin. To do so, we used a HEK293 cell line stably over-expressing APP^695WT^ (HEK293-APP^695WT^) and we observed a strong PLA signal (**Fig. 3F**). To validate the specificity of PLA signal, HEK293-APP^695WT^ were transfected with siRNA allowing the silencing of APP or talin. Both siRNA transfections resulted in a robust decrease in the number of PLA dots supporting the specificity of the PLA APP/talin signal (**Fig. 3G**). Next, PLA was performed on primary neuronal cultures to address the localisation of PLA APP-talin signal in neuronal context. For this purpose, PLA experiments were performed in association with standard immunofluorescence (IF) staining to visualise the synaptic marker, Homer. Co-localisation between PLA APP-talin and Homer staining supported the potential interaction of APP and talin at the synapse (**Fig.3H**).

### Structural modelling of APP as a synaptic adhesion molecule connecting the talin cytoskeletal connection to the synaptic junction

In integrins, NPxY motifs represent the foundations on which the cytoskeletal mechanosensory machinery assembles. Talin and kindlin bind to these anchor points and connect them to the actin and microtubule cytoskeletons. As shown in Fig.3, 4, <u>Movie 1</u> and <u>Movie 2</u>, four APP molecules that are dimerised in cis and trans clearly connect the two membranes together, precisely positioning two NPxY motifs in each of the pre– and post-synaptic compartments. In these figures for simplicity, we do not draw, or model, integrin or kindlin, even though it is likely that both are involved in the mechanocoupling of APP to the cells’ force generation machinery via talin (Fig.4).

Our analysis indicates a novel role for APP as a mechanocoupler, providing a physical means of synchronisation of the pre– and post-synaptic force generation machineries. In this arrangement, APP would be playing a central role in maintaining mechanical homeostasis, since forces generated across the synapse would be experienced by the APP mechanocoupling. This also indicates a mechanical basis for how and why APP can be differentially processed (and misprocessed) by being cleaved at subtly different positions. APP would be under varying amounts of tension and as such differential processing of APP would provide a mechanism for the cells to sense and respond accordingly to maintain mechanical homeostasis.

### Talin1 depletion dramatically alters APP processing in cells

Since synaptic dysfunction and loss is one of the very early hallmarks of AD correlating with earliest cognitive decline, we investigated the potential impact of talin1 expression level on APP processing. The processing of APP is complex and involves sequential enzymatic cleavage via enzymes referred to as secretases, that are located in both the interior and exterior compartments of the neuron (Wang et al., 2017). In addition, the identification of familial, AD-linked mutations in the genes for APP and presenilin (PSEN1 and PSEN2) associated with dysregulation of Aβ peptide production suggests that APP processing is at the heart of the disease process. In this context, intense research has focussed on exploring which proteins impact APP metabolism, and many proteins have been shown to alter the dynamic process of APP processing. We previously used a genome-wide, high-content siRNA screening (HCS) approach to measure the effect of depletion of each protein in the human proteome on APP metabolism (Chapuis et al., 2017). Talin1 and kindlin2 were among the top 5% of hits showing the strongest variations on APP metabolism (Fig.5A). This approach uses a rapid detection and quantification of intracellular APP fragments in HEK293 cells stably over-expressing a mCherry-APP^695WT^-YFP (Fig.5B). Talin1 silencing was associated with a significant increase in both mCherry and YFP fluorescence levels. Of note, the effects were similar to those observed after kindlin2 silencing (Chapuis et al., 2017).

**Figure 5.**
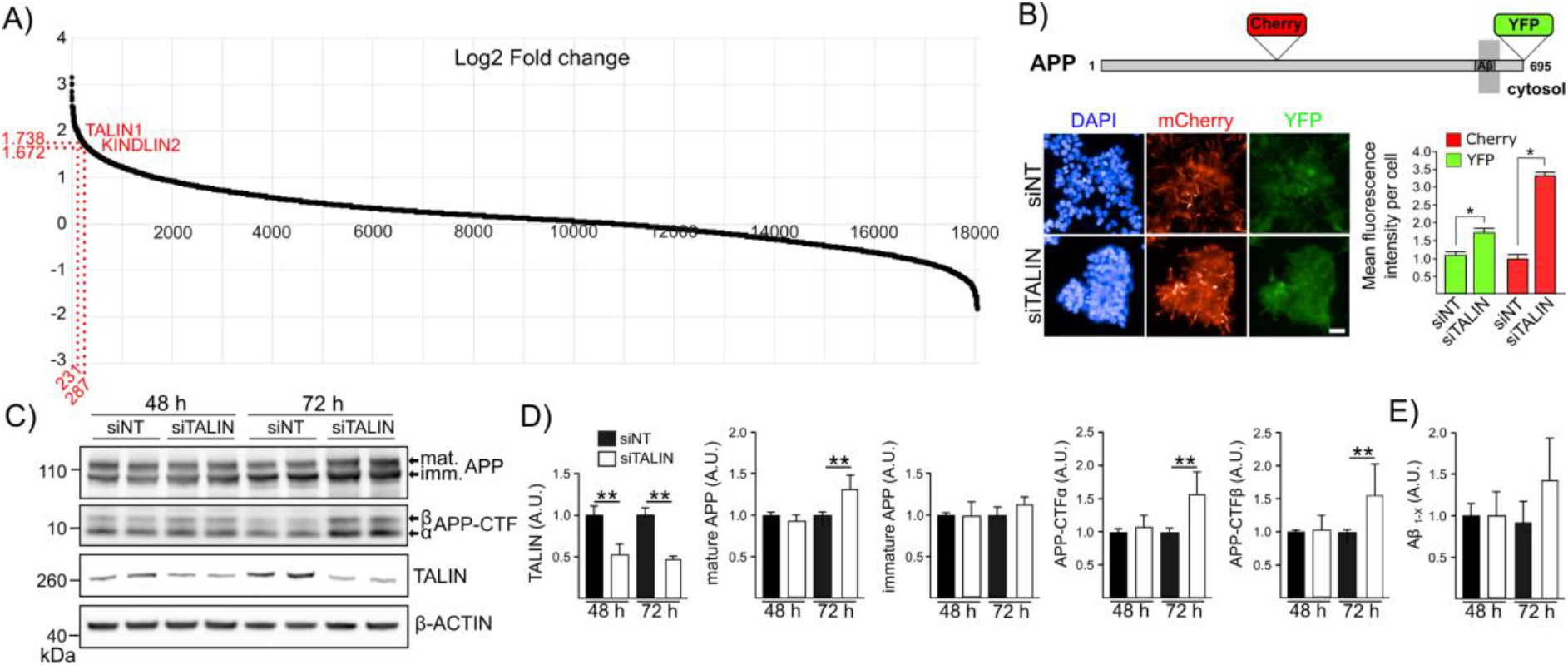
Talin1 depletion dramatically alters APP processing in cells. **A**) The published data set from (Chapuis et al., 2017), showing the distribution of modulation (Log2 Fold Change) of APP metabolism on genome-wide siRNA screening. The log2 fold change for talin1 (1.74) and kindlin2 (1.67) are shown. **B**) Schematic representation of APP, showing where the fluorescent proteins (mCherry and YFP) are inserted for the high-content siRNA screen (HCS). Representative fluorescence microscopy images and quantification showing the impact of talin1 silencing on mCherry and YFP intensity based on HCS data. Scale bar = 20 µm. **C-D) Impact of talin1 silencing on APP metabolism in the HEK293-APP695^WT^ cell line. C)** Cells transiently transfected with siTalin1 or non-targeting siRNA were analysed by Western Blot (WB) using anti-APP C-terminal, anti-talin1 or anti-actin antibodies. **D)** Densitometric analyses and WB quantifications from three independent experiments are shown. Mature APP, immature APP, C-terminal fragments (APP-CTF) α and β. **E)** Aβ_1-X_ secreted into conditioned medium were assayed using an AlphaLISA. Histograms indicate the mean ± SD. a.u., arbitrary units. ** p<0.01, non-parametric test.

Next, we investigated the impact of talin1 silencing on APP processing in HEK293-APP^695WT^ in absence of mCherry and YFP tags using conventional approach to quantify the main byproducts of APP. Upon talin1 silencing we observed a strong increase in mature APP levels and the accumulation of all the APP-derived substrates for α-, β– and γ-secretases (CTFα and CTFβ intracellular C-terminal fragments of APP produced respectively by α– and β-secretases, as well as Aβ secretions) (Fig.5C,D). Overall, our results support the involvement of talin expression level in the regulation of APP processing.

## Discussion

Here we present evidence of a direct coupling between APP and talin, identifying a new mechanism for connecting the mechanosensitive force generating machinery to the synaptic junction. In a correctly functioning connection, we propose that this coupling would serve as a force feedback mechanism that works to mechanically synchronise the two sides of the synapse ensuring mechanical homeostasis (Fig.6). The coupling between APP and the mechanical switches in talin offers several new testable hypotheses regarding the mechanical functioning of synapses and memory storage and how defects in this mechanocoupling can lead to AD. We present this discussion section as a series of six testable hypotheses that emerge from this new discovery.

**Figure 6.**
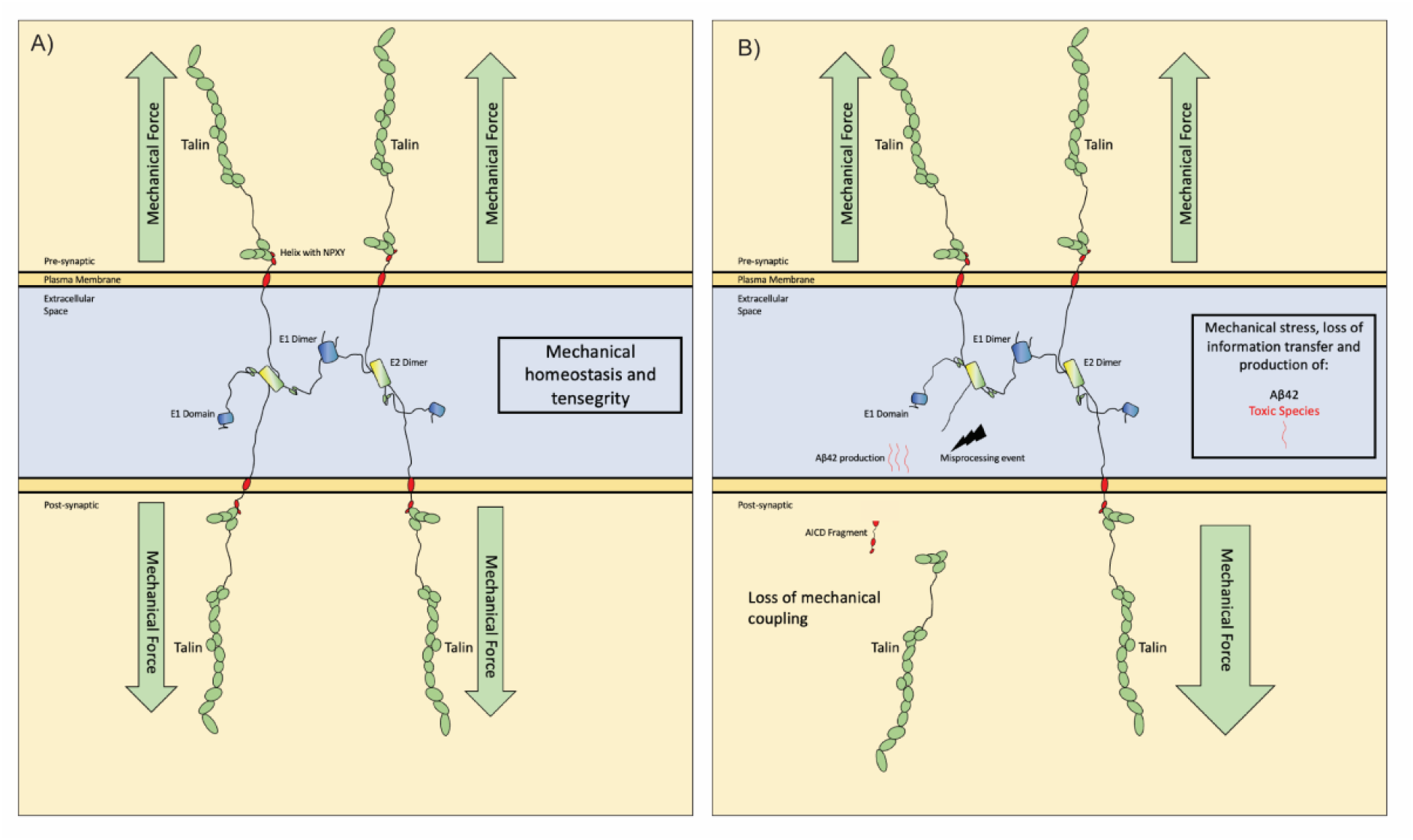
Concept model for the role of the APP-talin mediated mechanocoupling at the synapse in maintaining mechano-homeostasis, and how dyshomeostasis would lead to wholesale synaptic changes. **A**) In a healthy synapse the basal levels of mechanical forces across the synapse are balanced to establish mechanical homeostasis. Each synaptic stimulation event would lead to a transient increase in tension that would alter the switch patterns and update the synapse, before mechanical homeostasis is reestablished. APP, as part of the mechanocoupling of the synapse, would help to maintain the synchronisation of the system as altered mechanics or imbalance would lead to APP-processing driven feedback mechanisms that reinstate homeostasis. **B)** In AD this mechanical feedback mechanism would be defective, and misprocessing of APP would occur, accelerating the loss of mechanohomeostasis, leading to inappropriate force loading on the remaining cytosketal couplings and production of the toxic Aβ42 species. This loss of mechanical homeostasis would propagate through the two neurons spreading throughout the neuronal circuits as the entire systems mechanical synchronisation is lost.

**Figure 7.**
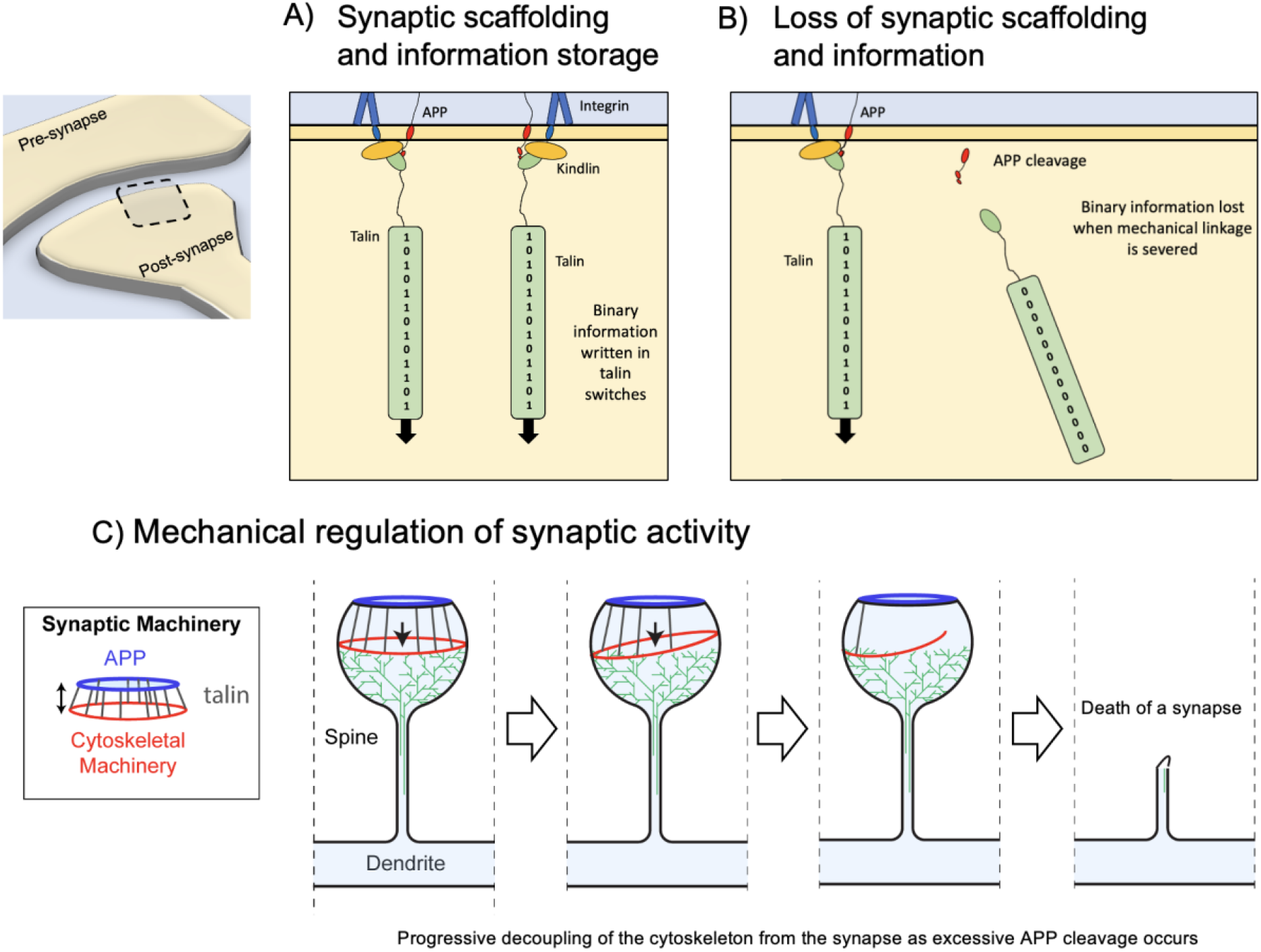
A concept model of mechano-homeostasis and dyshomeostasis leading to whole-scale synaptic changes. **A-B**) In the MeshCODE theory of a mechanical basis of memory, the synaptic scaffold protein talin is a memory molecule. The 13 force-dependent binary switches in the talin rod can store information in a binary format that spatially organises the synaptic enzymes to control synaptic activity. **A)** In a normal synapse the scaffolding controls the synaptic activity as a function of the talin switch patterns. The connection of talin to the membrane involves APP (red) and the Focal Adhesion complex containing integrin (blue) and kindlin (orange). **B)** Perturbed mechanical cues and destabilisation of the adhesion complex leads to increased APP processing and loss of the APP-talin connection. The information written in the shape of the talin molecule is lost as it is no longer under tension and the molecule resets. **C)** The coupling of the force generation machinery to the synaptic cleft provides a way to control the synaptic activity as a function of the switch patterns organising the synaptic enzymes differently. However, as each APP is cleaved, a connection of the cytoskeleton to the synaptic cleft is severed, leading to loss of information but also to dyshomeostasis. Ultimately the loss of mechanical couplings leads to the death of the synapse.

### Hypothesis 1: APP forms an extracellular meshwork that mechanically couples the two sides of the synapse

Drawing APP molecules to scale, and integrating all the known structural elements, provides a striking new view of APP’s role as a synaptic adhesion molecule that oligomerises to form an extracellular meshwork that directly couples the mechanosensitive machinery on both sides of the synapse (Fig.6). Our previous structural analysis of talin and vinculin in the synaptic scaffolds demonstrated the power of visualising the dimensions of synaptic proteins to scale (Barnett and Goult, 2022). AlphaFold structural models are very useful in this regard as they include the disordered regions of proteins that are often overlooked, thus enabling us to precisely model these complexes and the dimensions of the cytoskeletal linkages (Fig.4).

The mechanosensitive machinery of the cell is coupled to the plasma membrane via talin binding to the NPxY motif of integrins. The crystal structure of the NPxY motif of APP bound to talin identifies the APP cytoplasmic tail as a second attachment site for talin to engage, with the potential to forge similar cytoskeletal connections at the synaptic junction (Fig.6). APP has previously been shown to bind to kindlin2 (Dourlen et al., 2019; Eysert et al., 2021) and extracellularly to β1– and α3-integrins (Hoe et al., 2009; Young-Pearse et al., 2008). Both talin and kindlin2 are FERM domain-containing proteins and both bind to NPxY motifs on the integrin tail. Furthermore, talin also binds to kindlin2 (Aretz et al., 2023). So, there is substantial evidence of the role of APP in cell adhesion functions, but also of a role for cell adhesion proteins in the regulation of APP processing (Dourlen et al., 2019; Eysert et al., 2021). We propose that the tight interplay between cell adhesion and APP oligomerisation defines APP as a mechanocoupler, linking the cytoskeletal machineries on both sides of the synapse (Fig.6). Many FA proteins are genetic risk factors for APP processing. However, whilst talin is not in the GWAS for AD risk factors, many talin binding partners and FA regulatory components are (Dourlen et al., 2019; Eysert et al., 2021). It is likely that talin is less tolerant to variants as a result of its central role in mechanical computation in all cells.

### Hypothesis 2: APP processing is a mechanical signalling pathway that synchronises the synapse to ensure mechanical homeostasis

The formation of an APP-mediated mechanical coupling between the two sides of the synapse immediately indicates that APP processing will be affected by tension on APP, literally pulling/pushing the APP extracellular regions relative to the membrane-anchored secretases. In this way, we propose that APP’s role in neuronal function is to orchestrate a mechanically regulated signalling pathway, whereby altered tension on the synapse leads to differential APP processing and cellular signals on both sides of the synapse that re-establish mechanical homeostasis. Disruption of the tension on APP would provide a mechanism for the relative position shift that is key to APP misprocessing and toxic species formation.

The concept of mechanical regulation of proteolytic processing of signalling proteins is established in the case of Notch signalling (Bray, 2006). Notch serves as a mechanical signalling system, where protein cleavage is the signalling output (Guénette et al., 2017). The Notch ligands are typically membrane proteins on adjacent cells. In the case of APP, the ligands are other APP molecules on the adjacent cell at the synaptic junction. It is likely that APP and its homologs APLP1 and APLP2 represent a family of cell surface mechanical receptors. APP processing would therefore provide an essential signalling function in maintaining the mechanical integrity of the synapse. Thus, APP processing would ensure each synapse represents a perfect, isolated mechanical system which is essential if mechanical signals are to be transmitted with high fidelity as required for a mechanical basis of memory as proposed by the MeshCODE theory (Goult, 2021).

We show that talin depletion dramatically alters APP processing in cells (Fig.5) supporting the role of mechanical signals in APP processing. Previous experimental evidence in support of talin’s role in APP processing include; 1) APP immunoreactivity is co-localised with talin immunoreactivity in primary rat neuronal cultures, including hippocampal, cortical and cerebellar tissues, (Storey et al., 1996), 2) talin2 has been implicated in AD (Gusareva et al., 2018) and 3) talin has been linked to APP both in mouse models of AD (Lachén-Montes et al., 2016) and in platelets of AD patients (González-Sánchez et al., 2018).

### Hypothesis 3: A mechanical basis of AD – altered mechanical cues lead to misprocessing of APP which leads to the devastating consequences of AD

Visualising APP processing to scale (Fig. 6) led to our hypothesis that the mechanical couplings through the cytoskeletal-talin-APP connection are regulating the spatial positioning of the APP substrate relative to the secretase APP-processing machinery. This provides a way to mechanically synchronise the two boutons during development and healthy neuronal function, but excess or misprocessing would be deleterious. During synaptogenesis and synapse maintenance this coupling of mechanics and intracellular signalling would synchronise the two connecting cells and allow the precise mechanical couplings to form.

What is striking is how modest the changes are in processing between healthy and toxic states, a 40-residue fragment is relatively benign, but a 42-residue fragment is toxic. This subtle, 2 amino acid change in where γ-secretase cuts has huge ramifications for the system. As the β– and γ-secretases are positioned in the membrane, their active sites are relatively fixed and spatially restricted. We therefore propose a mechanical aspect to APP processing whereby tension on APP alters the position of the APP relative to the active site of the secretases leading to alternate processing. Altered mechanical inputs due to altered APP tension would be the cause of the two amino acid shift (7 Å) misprocessing.

### Hypothesis 4: Loss of memory in AD is a result of corruption of the binary coding within the synapses leading to loss of information

The importance of APP in the development of AD is well established and mis-regulated APP processing, leading to neuro-toxic fragments, is widely studied. However, the precise mechanisms behind the memory loss in AD are not fully understood, although death of synapses is clearly a factor (Selkoe and Hardy, 2016). Our analysis indicates that APP is critical for ensuring mechanical homeostasis, as its processing ensures the two connected neurons at a synapse are mechanically synchronised and able to transmit mechanical signals with high fidelity. This identifies a basis for a direct coupling of APP to the cell’s mechanical computation machinery. We previously proposed the MeshCODE theory of a mechanical basis of memory. This theory is based on the realisation that talin is comprised of binary switch domains that are operated by small changes in mechanical force. The 13 binary switches in talin present a way for information to be written into the synaptic scaffolds (Goult, 2021). If, across the entire synapse and the whole neural network, these structural states are forming a binary coding that encodes information in the form of memories, then mechanical dyshomeostasis leading to the MeshCODE being corrupted would be a consequence of perturbed APP processing.

This leads us to propose that the memory loss in AD comes as the information written in each synapse is lost and they start to desynchronise. AD leads not just to decoupling the synapse, at the synapse level but also relative to the circuit that synapse is part of. Amyloid plaques can appear up to 20 years before detectable cognitive defects (Selkoe and Hardy, 2016), which might indicate that there is considerable redundancy in memory storage structures in the brain. For a while the redundancy in memory storage would be able to compensate for this corruption of parts of the disk until the loss of information becomes overwhelming.

### Hypothesis 5: The spread of AD pathology is due to the collapse of mechanical homeostasis that propagates through the networks

What is striking about AD pathology is that it spreads from foci, and then propagates through the circuits across the brain. One question is why/how does it spread? Formation of Aβ42 or spread of tau seeds have been presented as possible answers to this question. Here we propose the alternative explanation that the spreading is a result of the slow collapse of the mechanical synchronisation of the brain.

When forces act on the synapse in healthy neurons, the APP-processing mechanical feedback systems described here will work to re-establish mechanical homeostasis. However, if the feedback system itself is defective then loss of homeostasis will be persistent and will impact on both connected neurons. If the patient is already primed towards AD (either by mutations in the secretases, or in the FA proteins that stabilise the synaptic junction along with APP), then any perturbation in mechanics will trigger improper APP processing at other synapses in the neurons. In this way, it would be the loss of mechanical homeostasis and the shockwave-like spread of it from the initial foci(s) that would be why AD spreads across the brain. This idea is supported by the existence of AD-causing presenilin mutations that counter-intuitively limit cleavage of APP and hence reduce APP processing (Bagaria et al., 2022; Sun et al., 2017). However, these mutations still give rise to AD, fitting with the idea that this is due to defective force feedback signalling, with the result being that they also accelerate mechanical dyshomeostasis. Functional spreading of AD, whereby as neurons become compromised they then compromise the integrity of neurons in adjoining areas has been reported (Khan et al., 2014) and the spread of mechanical dyshomeostasis identified here provides an explanation for this phenomenon.

### Hypothesis 6: It might be possible to repurpose drugs that stabilise Focal Adhesions to slow down the spread of AD

Hypothesis 5 indicates that if an individual is not pre-disposed to AD, then the robust APP-processing mechanical signalling pathway is able to maintain and re-establish mechanical homeostasis following each mechanical perturbation. However, disruption of this intimate coupling of the synapse, either via excess tension, mutations that destabilise this coupling, or other means would alter the availability of cleavage sites in APP leading to altered processing.

What is striking is that many of the GWAS risk factors identified for AD are in FA proteins or cytoskeletal regulators of FAs (Chapuis et al., 2017; Dourlen et al., 2019), indicating that a loss of adhesion integrity or stability contributes to AD progression. Similarly, APP knock down has been reported to impact on FA stability (Ristori et al., 2020) further supporting the idea that the stability and integrity of these adhesion complexes is critical for healthy brain function. The identified risk factors in GWAS include Src, Rac, Rho, kindlin2 and paxillin that are all known to modulate FA dynamics (Dourlen et al., 2019) and regulate FA stability. These mutations that impact on the mechanical attachments of the cytoskeleton to the synapse are genetically linked to AD, making it reasonable to assume that perturbations in the mechanics of the synaptic junctions are drivers of AD. It might be imagined that such destabilised FA^1^ connections lead to defective mechanics and as a result, aberrant APP-processing and reduced capacity for correction and re-establishment of homeostasis. Furthermore, on a systems level, if the cytoskeletal connections to the synapse are defective, then this would accelerate the spread of the dyshomeostasis, as the impaired connections would result in aberrant APP-processing at each synaptic connection as they fail to recover mechanical homeostasis.

This leads to Hypothesis 6, that therapeutic interventions that stabilise FAs in other cell types might represent a new approach to slowing the spread of AD. If destabilisation of FA dynamics in the synapse can be shown to be a driver of accelerated AD progression, it follows that stabilisation of FA via pharmaceutical means might present a novel therapeutic opportunity for slowing the spread of AD and thus delaying the onset of symptoms. Thus, we propose that it might be possible to repurpose already available, FDA approved drugs and pharmaceutical agents that stabilise FAs for use in AD to slow down the spread of AD through the brain. There are many such drugs already in existence that target effectors of FA dynamics. For example, i) focal adhesion kinase (FAK) inhibitors – FAK activity is shown to lead to adhesion turnover, so suppressing FAK would be one option, ii) Rho activators – increase activation to increase actomyosin contractility which has been shown to stabilise FAs, iii) integrin activators – many compounds and antibody strategies exist for enhancing integrin activation, in doing so these stabilise FAs. iv) microtubule disruptors – in cultured cells, microtubule targeting of FAs can lead to the turnover of FAs, therefore reducing microtubule-dependent FA turnover would stabilise FAs.

## Summary

Here we present 6 novel hypotheses for the role of Amyloid Precursor Protein (APP) in healthy neuronal activity and its misprocessing and memory loss in Alzheimer’s Disease.

## Hypotheses

**Hypothesis 1:** APP forms an extracellular meshwork that mechanically couples the two sides of the synapse.

**Hypothesis 2:** APP processing is a mechanical signalling pathway that synchronises the synapse to ensure mechanical homeostasis.

**Hypothesis 3:** A mechanical basis of AD – altered mechanical cues lead to misprocessing of APP which leads to the devastating consequences of AD.

**Hypothesis 4:** Loss of memory is a result of corruption of the binary coding within the synapse.

**Hypothesis 5:** The spread of AD is due to the collapse of mechanical homeostasis that propagates through the networks.

**Hypothesis 6:** It might be possible to repurpose drugs that help stabilise Focal Adhesions to slow down the spread of AD

Finally, we note two additional points; i) APP expression is not exclusive to neurons and is found in all cells and ii) in evolutionary terms, APP-like proteins preceded synapses by millions of years (as did talin). As such, the coupling of APP processing to the mechanical computation machinery we identify here may be playing a more global role in maintaining mechanical synchronisation of cells and possibly represents a force feedback signalling mechanism in all animal cells.

## Limitations of this study

Here we present evidence of a direct link between talin and APP which points to a completely new view of AD and the role of APP as a signalling pathway. In biochemical assays in solution the interaction is relatively weak, however it is specific and can be perturbed by targeted mutations. With the multivalency and contributions from the membrane and numerous other factors this interaction, the linkage will be much tighter in cells, as it is for the talin-integrin connection. It might also be that talin-kindlin-integrin-APP represents the stable complex.

## Materials and Methods

### DNA constructs

Talin1 F3, talin1 F2F3, talin2 F3 and talin2 F2F3 constructs were as described previously (Anthis et al., 2009). The chimera was designed based upon the strategy and crystallised structure of integrin β3-talin (PDB ID: 1MK7) (Garcıá –Alvarez et al., 2003) in which the intracellular domain of APP (QNG**YENPTY**KF) was fused to the N-terminus of talin1 F2F3. The APP-talin chimeric construct fused APP residues 754-764 with talin residues 209-400 to generate an APP(754-764)-talin(209-400) chimera. This sequence was ordered from GeneArt as a codon-optimized synthetic gene (APP-F2F3) in the pet151 expression vector.

### Protein expression and purification

Talin1 F3, talin1 F2F3, talin2 F3, talin2 F2F3 and APP-F2F3 chimera constructs were expressed in BL21(DE3) *E. coli* cells. Cells were harvested and stored at –80°C in lysis buffer (50 mM Tris pH 8.0, 250 mM NaCl). Proteins were purified using previously described methods (Khan et al., 2021). Succinctly, the cells were lysed by sonication and the soluble fraction was loaded onto a 5 mL HisTrap HP column (Cytiva), then eluted across a 15-column volume (CV) linear imidazole gradient. Protein was dialysed overnight in 20 mM sodium phosphate pH 6.5, 50 mM NaCl with TEV protease (AcTEV; Thermo Fisher) added to remove the His-tag. All proteins except talin2 F3 were further purified by cation-exchange chromatography using a 5 mL HiTrap SP HP column (Cytiva) and eluted across a 15 CV linear NaCl gradient. Talin2 F3 was further purified by anion-exchange using a 5 mL HiTrap Q column. Proteins were dialysed against PBS pH 7.4, concentrated, flash frozen in LN_2_ and stored at –80°C.

### Nuclear Magnetic Resonance (NMR) binding studies

For NMR binding studies, cells were grown in ^15^N-labelled 2M9 minimal media (850 mL Milli-Q water, 100 mL 10x M9 salts, 1 mL 0.1 M CaCl_2,_ 1 mL of 1 M MgSO_4_, 10 mL BME vitamin solution (Sigma-Aldrich), 4 g glucose, 1 g ^15^N-labelled ammonium chloride per litre and 100 µg/mL ampicillin). ^15^N-labelled protein samples were prepared at 150 µM final concentration in 20 mM phosphate buffer pH 6.5, 50 mM NaCl, 2 mM DTT, 5% (v/v) D_2_O. NMR spectra were collected at 298 K on a Bruker Avance III 600 MHz NMR spectrometer equipped with Cryoprobe. All data were processed using TopSpin and analysed with CCPN Analysis (Skinner et al., 2015).

### Fluorescence Polarisation Assay

Peptides containing a non-native N-terminal cysteine were synthesised by GLBiochem (Shanghai): APP(732-770) C-HHGVVEVDAAVTPEERHLSKMQQNGYENPTYKFFEQMQN, APP(732-770)-4A C-HHGVVEVDAAVTPEERHLSKMQQNGYEAAAAKFFEQMQN. Integrin Beta1A(752-798) CKLLMIIHDRREFAKFEKEKMNAKWDTGENPIYKSAVTTVVNPKYEGK and KANK1-(30-60)4A PYFVETPYGFQAAAAFVKYVDDIQKGNTIKKLNIQKRRKC peptides were as described previously (Bouchet et al., 2016). Peptides were labelled with a maleimide-fluorescein dye (Thermo Fisher) following the manufacturer’s protocol.

Assays were performed in PBS pH 7.4, 0.01% v/v Tween-20, in triplicate with 500 nM peptide and a 2-fold serial dilution of protein. Fluorescence polarisation was measured using a CLARIOstar plate reader (BMG LABTECH) at 25°C (excitation: 482 ± 8 nm; emission: 530 ± 20 nm). Data were analysed using GraphPad Prism 8 software and K_d_ values were generated using a single-site total binding model.

### Crystallization, X-Ray Crystallography and Structural Determination

The APP-talin1(F2F3) chimera was crystallised using hanging-drop vapour diffusion. Sparse-matrix crystallisation trials were set up using a Mosquito LCP (TTP LabTech) with 100 nL protein at 10 mg/mL mixed with 100 nL mother liquor and stored at 20°C. Microcrystals formed in 0.1 M Tris pH 8.5, 20% v/v ethanol, which were manually optimised for pH, ethanol and protein concentration in 2 µL hanging drops with a 1:1 protein to mother liquor ratio. Crystals formed at 6 mg/ml protein in 0.1 M Tris pH 8.5, 10% v/v ethanol at 18°C and were harvested by transferring the crystal to fresh crystallisation solution supplemented with 40% v/v glycerol and cryocooled in LN_2_.

The crystals belong to the space group P2_1_2_1_2_1_ with a single copy of the APP-talin1(F2F3) chimera in the asymmetric unit (AU). X-ray diffraction data were collected at 100 K on the I04 beamline at Diamond Light Source (Didcot, UK). The data were moderately anisotropic and were processed using Xia2/DIALS (Beilsten-Edmands et al., 2020; Winn et al., 2011; Winter, 2010; Winter et al., 2018). The structure was solved by molecular replacement with PHASER (McCoy et al., 2007) using the talin(F2F3) domains as a search model (1MIX; (Garcıá –Alvarez et al., 2003)). After initial placement, there was obvious difference density at the N-terminus corresponding to the APP. The model was completed through iterative rounds of model building in COOT (Emsley et al., 2010) and reciprocal space refinement using PHENIX (Afonine et al., 2012; Liebschner et al., 2019).

The final model comprises residues 209-400 of talin1(F2F3), with residues K322 and N323 missing from the loop in F3 that connects β1 and β2. The mainchain density for the APP (754-764) is well defined and the side chains for N759 – F764 (including the NPxY motif) could be unambiguously placed. The model was refined to an *R_work_*/*R_free_* of 21.1/26.7 and has good geometry as determined by MolProbity (Williams et al., 2018), with 95% of residues in the preferred region of the Ramachandran plot, 5% in the additionally allowed region and a single outlier. Details of the crystal parameters, data collection, processing and refinement statistics are shown in Supplementary Table 1. The coordinates and structure factors were deposited in the PDB with accession code 8S4Y.

### Structural Modelling

The predicted AlphaFold 2.0 structural models of APP770 (UniProt ID: P05067-1), APP751 (UniProt ID: P05067-4), APP695 (UniProt: P05067-8) were obtained from the AlphaFold Protein Structure Database (Jumper et al., 2021; Varadi et al., 2021). The atomic structures used in this study were;

3NYL: Crystal Structure of the E2 dimer of APP (Wang and Ha, 2004)

3KTM: Crystal structure of the heparin-induced E1 dimer of APP (Dahms et al., 2010)

4JFN: Crystal structure of the N-terminal, growth factor-like domain of the amyloid precursor protein bound to copper (Baumkötter et al., 2014)

6R9T: CryoEM structure of full-length talin1 (Dedden et al., 2019)

To visualize the APP structures in the context of a plasma membrane, we used a previously modelled DOPC (1,2-Dioleoyl-sn-glycero-3-phosphocholine) lipid membrane PDB file (Kukol, 2009).

### Primary neuronal cultures

Animal housing and experimentation were performed according to procedures approved by the local Animal Ethical Committee following European standards for the care and use of laboratory animal (agreement APAFIS #32824-2021120518521661, Lille, France). Culture media and supplements were from Thermo Fisher, unless otherwise stated. Cortical neurons were dissected from E14-E15 mice. Briefly, cortices were isolated from E14-E15 mice in ice-cold dissection medium (Hank’s balanced salt solution supplemented with 10 mM HEPES, 1 mM sodium pyruvate, 10 mM glucose, and penicillin/streptomycin) and trypsinized at 37°C for 30 min (Trypsin solution, T4549, Sigma). DNase I was added to the trypsin-incubated tissue suspension to break down DNA and to avoid clumping of tissue during the subsequent trituration (DN25, Sigma). Trypsin was inactivated by the addition of isolation medium (Neurobasal™ medium supplemented with 10% heat-inactivated fetal bovine serum, 1% GlutaMAX, 20mM Hepes and Gentamycin). The cell suspension was passed through a 100 µm then 70 µm cell strainer followed by two centrifugations (0.3×g for 10 min). Cells were resuspended in culture medium composed of MACS Neuro Medium (130-093-570) supplemented with 0.25% GlutaMAX, 2% MACS NeuroBrew-21 (130-093-566) and Gentamycin and counted.

### Synaptosome extraction

To verify the presence of proteins at the synaptic level we did a subcellular fractionation as previously described (Eysert et al., 2021). Briefly, cortical neurons were resuspended in 0.32 M sucrose and 10 mM HEPES, pH 7.4 and centrifuged at 1,000× g for 10 min to remove nuclei and debris. The supernatant was centrifuged at 12,000× g for 20 min to remove the cytosolic fraction. The pellet was resuspended in 4 mM HEPES, 1 mM EDTA, pH 7.4 and was centrifuged 2× at 12,000× g for 20 min. The new pellet was resuspended in 20 mM HEPES, 100 mM NaCl, 0.5% Triton X-100, pH 7.2 for 1 h at 4°C and centrifuged at 12,000× g for 20 min. The collected supernatant corresponds to the non-PSD fraction (Triton-soluble). The remaining pellet was resuspended in 20 mM HEPES, 0.15 mM NaCl, 1% Triton X-100, 1% deoxycholic acid, 1% SDS, pH 7.5 for 1 h at 4°C and centrifuged at 10,000× g for 15 min to obtain a supernatant containing the PSD fraction (Triton-insoluble). The different fractions were then analysed by WB.

### Cell culture and transfection

Human embryonic kidney (HEK) 293 cells were cultured in 1:1 mixture of Dulbecco’s Modified Eagle Medium and Ham’s F12 nutrient mixture (DMEM-F12, 21331020, Thermo Fisher Scientific) supplemented with 10% heat-inactivated fetal bovine serum (FBS; 10270106, Gibco), 2 mM L-glutamine (25030149, Thermo Fisher) and 50 UI/mL penicillin/streptomycin (15140122, Thermo Fisher) at 37°C in a humidified atmosphere with 5% CO_2_. Prior to transfection, cells were plated at a density of ∼70%. For siRNA transfection, we used Dharmacon siRNA, non-targeting (D0018100105) and siTalin1 (L-012949-00-0005) and the Lipofectamine™ RNAiMAX (13778150, Thermo Fisher Scientific) was used as transfection reagent according to the manufacturer’s instructions. Cell line was tested negative for mycoplasma contamination using PCR test (Venor GeM OneStep, Minerva Biolabs).

### Immunofluorescence

Cells were fixed in 4% paraformaldehyde (PFA) for 15 min, washed 3× with PBS, and permeabilized for 5 min with 0.3% Triton X-100. HEK293-APP cells were incubated with 5% normal donkey serum for 1 h at RT before overnight incubation with the following primary antibodies: Talin1 clone 97H6 (1/400; NBP2-50320), anti-Amyloid precursor protein C-Terminal (1/500; A8717). The cells were then washed 3× with PBS and incubated with the following secondary antibodies raised in donkey (AlexaFluor-conjugated AffiniPure Fragment 488 or 647, Jackson ImmunoResearch) and 1/10,000 Hoechst 33342. Alternatively, for HEK293-APP cells and mouse embryo neurons, talin1 clone 97H6 (1/400; NBP2-50320), anti-Amyloid precursor protein C-Terminal (1/500; A8717), Homer1 (1/400; 160004) antibodies were used for the proximity-ligation assay (PLA) according to the manufacturer’s instructions (Duolink®, Olink Bioscience).

### Proximity Ligation Assay (PLA)

Cells were fixed in 4% paraformaldehyde (PFA) for 15 min, washed 3× with PBS, and permeabilized for 5 min with 0.3% Triton X-100. The proximity-ligation assay (PLA) was performed according to the manufacturer’s instructions (Duolink^®^, Olink Bioscience). Cells were incubated with the Duolink^®^ Blocking Buffer for 1 h at 37°C before overnight incubation with the following primary antibodies: Talin1 clone 97H6 (1/400; NBP2-50320), anti-Amyloid precursor protein C-Terminal (1/500; A8717). For mouse embryo neurons, Homer1 (1/400; 160004) antibody was also added. For the counterstaining, neurons were incubated with the following secondary antibodies raised in donkey (AlexaFluor-conjugated AffiniPure Fragment 405, 488 or 647, Jackson ImmunoResearch) and HEK293-APP were incubated with 1/10,000 Hoechst 33342.

### Western blot and Aβ quantification

Solutions and buffers were from Thermo Fisher, unless mentioned otherwise. Protein lysates were harvested in minimum volume of 100 µL/well in 24-well plates, in ice-cold lysis buffer as described previously (Chapuis et al., 2017) and quantified by means of Pierce™ BCA Protein Assay Kit (23225). Lysates were mixed with 1× lithium dodecyl sulphate (LDS) sample buffer (NP0008) and 1× reducing agent (NP0009), sonicated and boiled at 95°C for 5 min. 7 μg of total protein per well were loaded onto precast 4–12% Bis-Tris Protein Gels, and electrophoresis was achieved by applying 150 V for 90 min using an Invitrogen™ XCell SureLock™ Electrophoresis system with the NuPAGE® MOPS SDS running buffer (1×; NP000102). Proteins were transferred to a nitrocellulose membrane of 0.2 μM pore size (Bio-Rad) using the Trans-Blot Turbo Transfer System (Bio-Rad). Membranes were rinsed three times for 5 min in TNT (0.01 M Tris pH 8.0, 0.15 M NaCl, 0.05% Tween-20) and incubated with the following primary antibodies in SuperBlock T20 blocking buffer (Thermo Scientific) at 4°C overnight: mouse anti-talin1 (NBP2-50320; 1/1000 Novus Biologicals), rabbit anti-APP (A8717; 1/5000; Sigma) and mouse anti-β-actin (A1978; 1/10000; Sigma). Membranes were rinsed 3× for 5 min with TNT and then incubated with horseradish peroxidase (HRP)-conjugated secondary antibodies (HRP-anti-mouse and HRP-anti-rabbit; 1:10000; Jackson ImmunoResearch) diluted in 5% milk for 2 h at room temperature. Blots were developed using Amersham ECL Western Blotting Detection Kit (WBLUC0500; Millipore). β-Actin was used as loading control.

## Acknowledgements

C.E. and N.L.W were supported by PhD studentships from the Biological Sciences Research Council (BBSRC) SoCoBio-DTP (BB/T008768/1). B.T.G. was supported by Cancer Research UK Program Grant (CRUK-A21671). C.N. was supported by France Alzheimer Program Grant (Genetic of Synaptic Dysfunction, 6362, 2023). We thank Guillaume Jacquemet, Yasumi Otani and Katy Goult for critical reading of the manuscript.

## Supplementary Information

**Table 1.** Data collection and refinement statistics of the APP-Talin1(F2F3) chimera (PDB ID:8S4Y)

| **Data collection** |  |
| --- | --- |
| Space group | P2<sub>1</sub>2<sub>1</sub>2<sub>1</sub> |
| Wavelength (Å) | 0.95375 |
| Cell dimensions |  |
| *a*, *b*, *c* (Å) | 60.39, 64.08, 65.19 |
| $\alpha$, $\beta$, $\gamma$ (°) | 90, 90, 90 |
| Resolution (Å) | 45.74 – 2.70 (2.74 – 2.70) |
| *R*<sub>meas</sub> (%) | 12.8 (85.0) |
| Total reflections | 47,956 (2,353) |
| Unique reflections | 7,374 (359) |
| *I* / $\sigma$(*I*) | 12.8 (1.9) |
| CC(1/2) | 0.993 (0.832) |
| Completeness (%) | 100 (97.0) |
| Multiplicity | 6.5 (6.6) |
| **Refinement** |  |
| Resolution (Å) | 45.74 – 2.70 |
| No. reflections (test set) | 7,322 (729) |
| *R*<sub>work</sub> / *R*<sub>free</sub> (%) | 21.1 / 26.7 |
| No. atoms |  |
| Protein | 1,714 |
| Water | 8 |
| *B*-factors |  |
| Wilson | 57.4 |
| Average | 66.0 |
| Protein | 67.2 |
| Water | 56.9 |
| R.M.S deviations |  |
| Bond lengths (Å) | 0.009 |
| Bond angles (°) | 1.032 |

*R*<sub>free</sub> is calculated using 10% of data isolated from refinement. Values in parentheses are for highest resolution shell.

**Figure S1.**
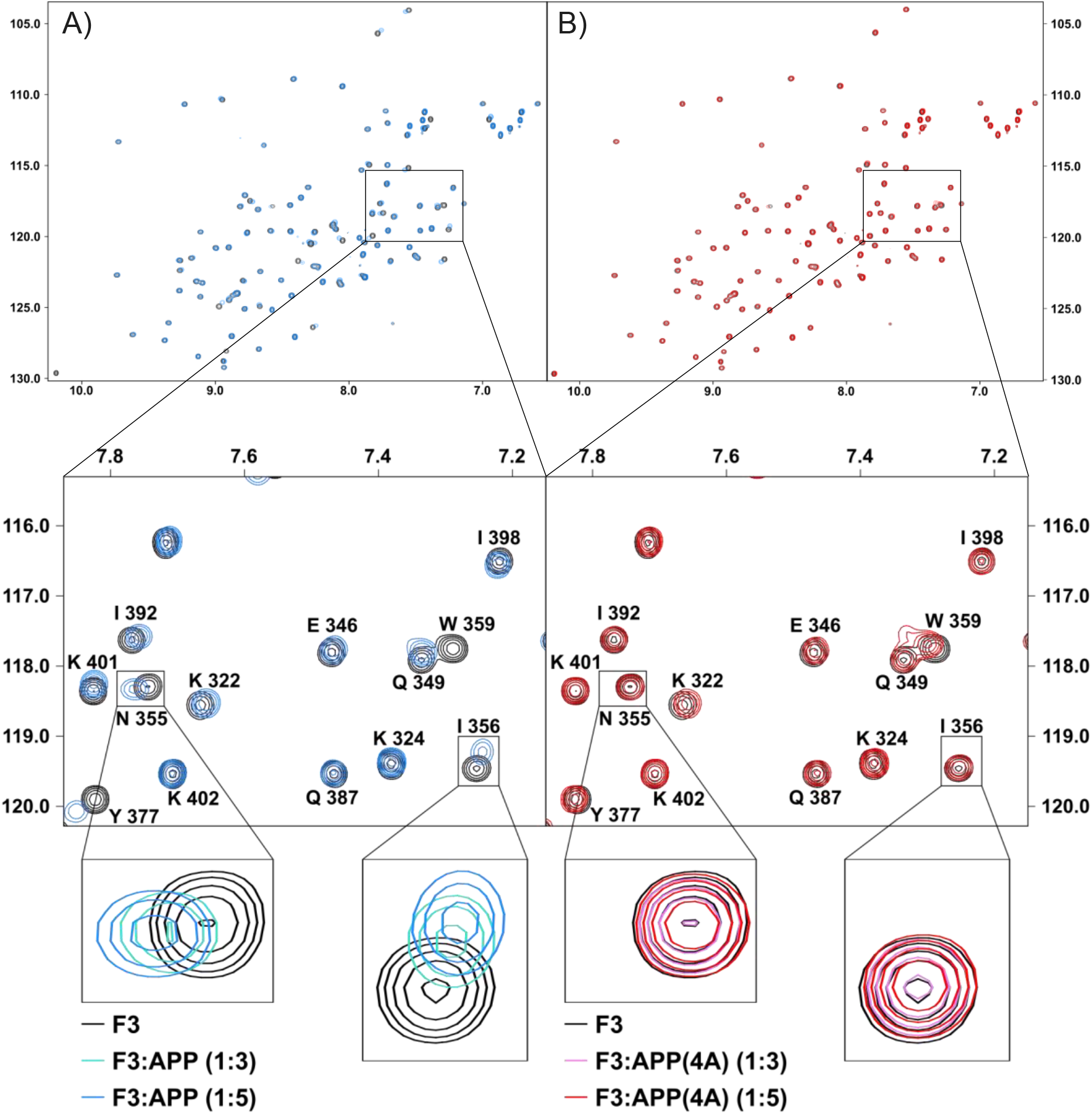
NMR spectra of the interaction of APP with talin1 F3 domain. 150 µM talin1 F3 on its own (black) and in the presence of **A)** wildtype APP (blue) and **B)** APP(4A) where the four residues of the NPTY motif are mutated to alanine (red).

**Figure S2.**
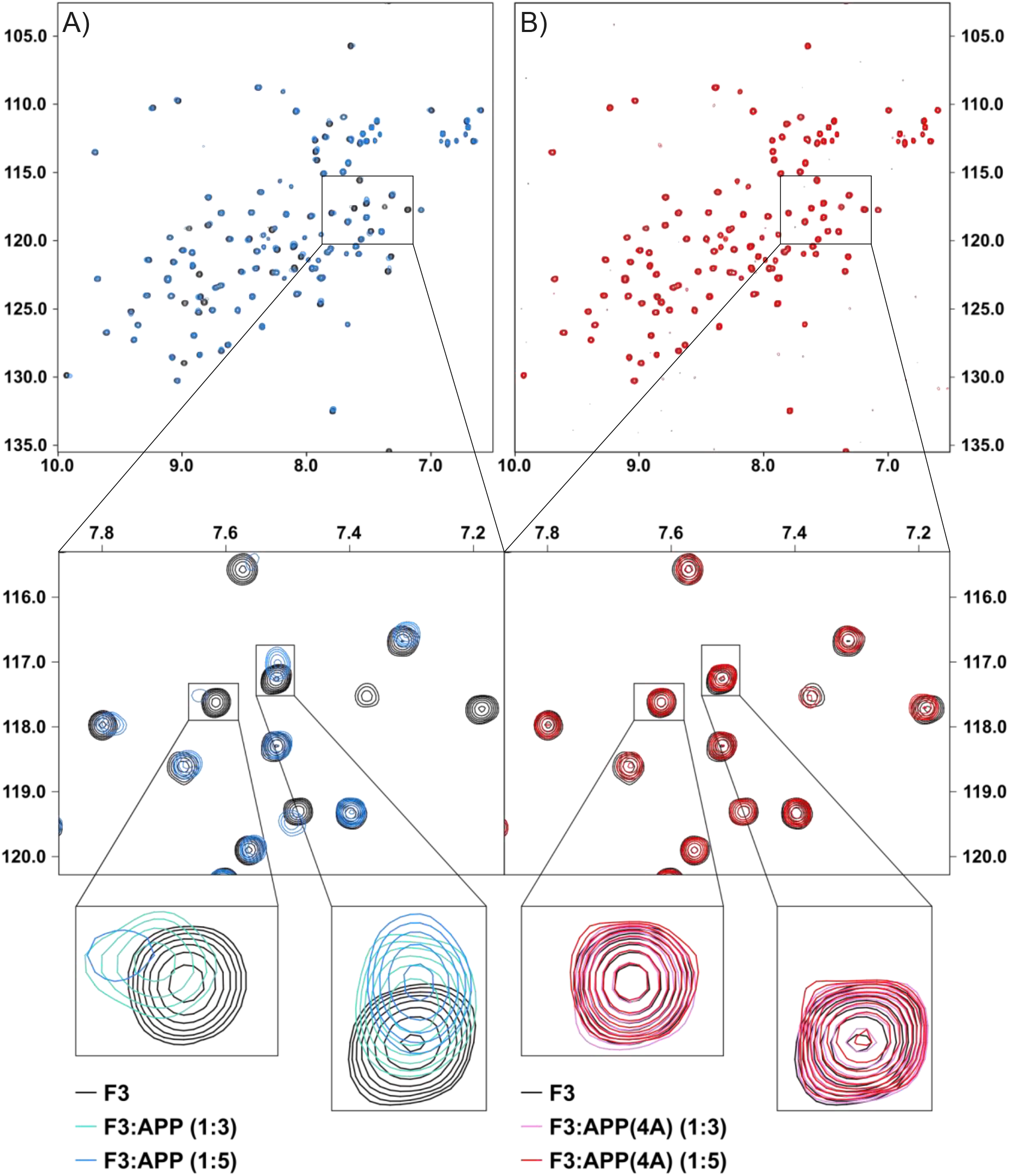
NMR spectra of the interaction of APP with talin2 F3 domain. 90 µM talin2 F3 on its own (black) and in the presence of **A)** wildtype APP (blue) and **B)** APP(4A) where the four residues of the NPTY motif are mutated to alanine (red).

**Figure S3.**
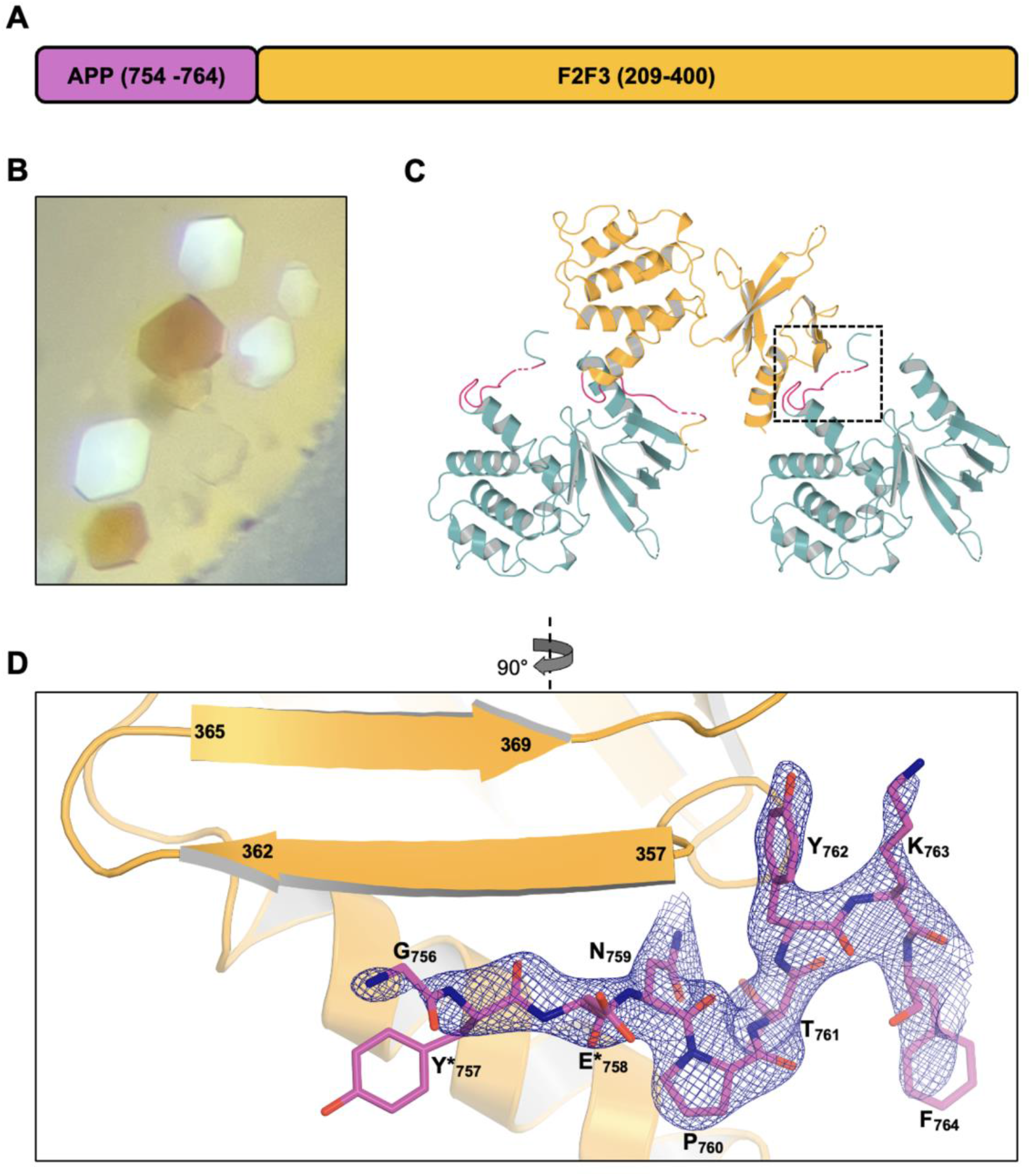
APP-F2F3 chimera design, crystallisation and structure. In all panels APP is coloured violet and F2F3 is coloured orange. **A)** Schematic representation of the APP-F2F3 chimera with the residues of APP shown in violet and the residues of F2F3 shown in orange. **B)** The chimera crystallised as birefringent flat plates. **C)** The chimera crystallised with a single copy in the asymmetric unit and the APP interacts with the F3 domain of a symmetry related molecule (teal) in a “daisy-chain” arrangement. **D)** Detail of the APP-F3 interaction (the boxed region in C)). The NPTY motif was unambiguously built into the electron density (2FoFc map contoured at 1.6 σ). The peptide backbone of residues G756-E758 is well defined and extends the β-sheet of the F3. Side chains not visible in the electron density (*) were set to zero-occupancy.

**Figure S4.**
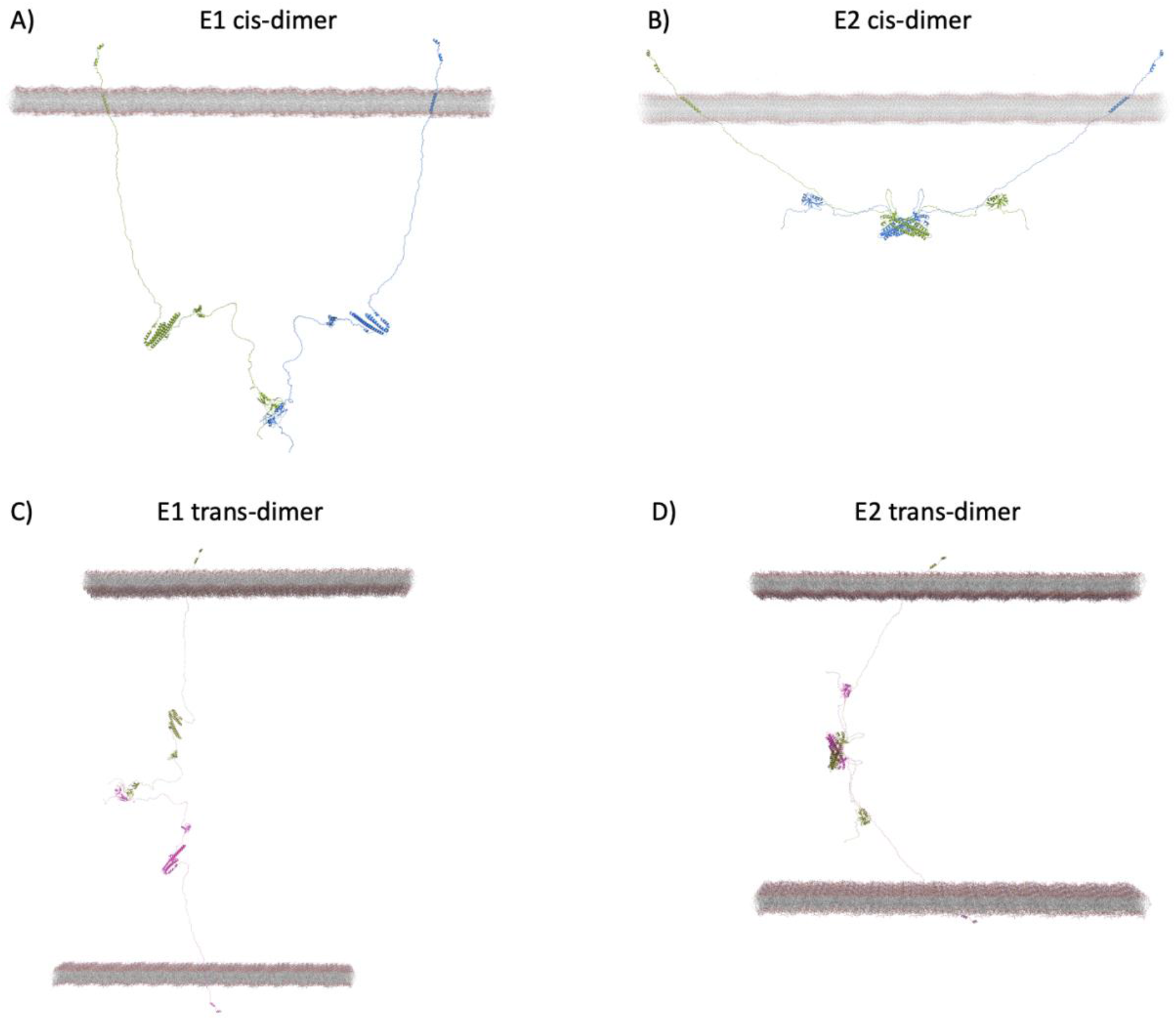
APP dimers in cis and trans. (**A-B**) Two configurations of an APP dimer in cis. **A)** APP cis-dimer mediated via the E1 domains. **B)** APP cis-dimer mediated via the E2 domains **(C-D)** two configurations of an APP dimer in trans. **C)** APP trans-dimer mediated via the E1 domains. **D)** APP trans-dimer mediated via the E2 domains.

## Movies

Movie 1. **Movie of the structural modelling of APP as a synaptic adhesion molecule connecting the talin cytoskeletal connection to the synaptic junction.** The movie is available at https://www.youtube.com/watch?v=csMs8-ZUHt4

Movie 2. **Movie summarising the structural modelling process in this study that revealed how APP can provide a mechanical linkage between the two neurons at a synapse.** The movie accompanies Figure 3. The movie is available at https://www.youtube.com/watch?v=qpz0wIpTlHM

## Footnotes

1 We note that technically the integrin adhesion complexes in neurons are not strictly Focal Adhesion’s. The integrin adhesion complexes in synapses are smaller and have been referred to as “Point contact” adhesions (Arregui et al., 1994; Nichol et al., 2016).

